# Neuropsychological Test Performance of Cognitively Healthy Centenarians: Normative data from the Dutch 100-plus Study

**DOI:** 10.1101/366328

**Authors:** Nina Beker, Sietske A.M. Sikkes, Marc Hulsman, Ben Schmand, Philip Scheltens, Henne Holstege

## Abstract

**Background:** The population who reaches the extreme age of 100 years is growing. At this age, dementia incidence is high and cognitive functioning is variable and influenced by sensory impairments. Appropriate cognitive testing requires normative data generated specifically for this group. Currently, these are lacking. We set out to generate norms for neuropsychological tests in cognitively healthy centenarians while taking sensory impairments into account.

**Methods:** We included 235 centenarians (71.5% female) from the 100-plus Study, who self-reported to be cognitively healthy, which was confirmed by an informant and a trained researcher. Normative data were generated for 15 tests that evaluate global cognition, pre-morbid intelligence, attention, language, memory, executive and visuo-spatial functions by multiple linear regressions and/or percentiles. Centenarians with vision and/or hearing impairments were excluded for tests that required these faculties.

**Results:** Subjects scored on average 25.6±3.1 (range 17-30, interquartile-range 24-28) points on the MMSE. Vision problems and fatigue often complicated the ability to complete tests, and these problems explained 41% and 22% of the missing test scores respectively, whereas hearing problems (4%) and task incomprehension (6%) only rarely did. Sex and age showed a limited association with test performance, whereas educational level was associated with performance on the majority of the tests.

**Conclusions:** Normative data for the centenarian population is provided, while taking age-related sensory impairments into consideration. Results indicate that, next to vision impairments, fatigue and education level should be taken into account when assessing cognitive functioning in centenarians.

## INTRODUCTION

Worldwide, the number of centenarians is expected to increase almost 20-fold to 3.2 million people in the next 30 years [1]. The incidence of dementia increases exponentially with age, reaching ~40% at age 100 [2]. Previous studies estimated that 25% of centenarians show no objective evidence of memory deficits, while 25% have symptoms of cognitive impairment and 50% may be regarded as having dementia [3, 4]. To evaluate cognitive impairment in this heterogeneous group, it is of great importance that suitable instruments be implemented that take into account the specific characteristics that are observed in centenarians.

Most commonly, evaluation of cognitive functioning in centenarians is based on normative data generated in younger elderly. However, the use of cut-off scores derived in younger samples may not account for the cognitive decline as part of the normal ageing process. Therefore, when these norms are used to assess cognitive performance of a centenarian, this may lead to misclassifications of cognitive impairment [5]. Several studies showed that specific cognitive functions decline during the process of ageing, including a slower processing speed and a decline in executive functions, whereas domains such as language and semantic knowledge are relatively maintained [6-8]. Between elderly aged 80 and 100 years, significant differences in cognitive test performance across cognitive domains were observed with increasing age [5, 9, 10]. This effect culminated in centenarians; their mean performance on cognitive test performance was lowest, while the variability in test performance was increased compared to 80 and 90 year olds [11, 12]. These findings suggest that accurate assessments of cognitive functioning of elderly at extreme ages can only be obtained using normative data generated in cohorts with relatively narrow age bands [9].

Thus far, most studies that provided normative data for centenarians described their performance on the Mini-Mental State Examination (MMSE) [12-16]. The MMSE is used to screen global cognitive functioning, while additional cognitive tests are required to evaluate a broad spectrum of cognitive domains. The Georgia Centenarian Study (GCS) provided data on tests that evaluate multiple cognitive domains, among which the Controlled Oral Word Association Test, the Fuld Object Memory Evaluation and Severe Impairment Battery and the Behavioral Dyscontrol Scale [12, 17-19]. However, these data (i) were generated in population-based samples, which included centenarians with cognitive impairment and (ii) were not adjusted for loss of hearing and sight. This will lead to lower norm-ranges, which might complicate the evaluation of cognitive health in a clinical setting. To evaluate cognitive performance in centenarians, it is necessary to compare their test performance with the performance of a large cognitively intact sample, and to take into account vision and hearing disabilities. The 90+ Study previously included non-demented subjects in the age bands (90-91, 92-94, 95+) to provide suitable normative data [10]. Despite the test adaptations made to compensate for sensory losses in the 90+ Study, some subjects could not complete tests due to sensory impairments, which emphasizes the importance of taking these faculties into account when establishing normative data.

Here, we aim to generate normative data for the evaluation of cognitive functioning in centenarians. We used a large sample of Dutch cognitively healthy centenarians from the 100-plus Study [20]. Per test, we considered whether vision and hearing abilities should be taken into account. The cognitive tests used in this study cover a wide range of cognitive domains including global cognition, pre-morbid intelligence, attention, language, memory, executive and visuo-spatial functioning.

## METHODS

### Subjects

Subjects were part of the Dutch 100-plus Study cohort, a longitudinal cohort study of people who 1) were aged ≥100 years, 2) self-reported to be cognitively healthy, which was confirmed by family members and/or caregivers. For this study, we specifically selected subjects who were estimated to be cognitively healthy by a trained researcher and of whom we had at least one neuropsychological test available. We excluded subjects with vision and/or hearing difficulties for tests requiring these sensory functions, leading to a total sample of N=235. A description of inclusion procedures can be found elsewhere [20]. This study was approved by the Medical Ethics Committee and all subjects provided informed consent.

### Procedure

#### Estimation of cognitive health, hearing and vision

The subjects were visited at home by researchers with a neuropsychological and/or medical training. The researchers noted their clinical impression of the cognitive health of the centenarians. This estimation was based on features such as centenarians continually repeating themselves, having difficulty understanding or remembering questions, and with naming and/or word finding. During study visits, vision and hearing were subjectively evaluated by the researcher and categorized into “good”, “moderate”, “poor” and “very poor” based on the features listed in the supplemental material. Subjects with poor to very poor vision were excluded from reporting normative data of the Mini-Mental State Examination (MMSE), the Key Search test, the Dutch Adult Reading Test (DART), the Visual Association Test (VAT), Trail Making Test (TMT), Number Location and the Clock Drawing Test (CDT). Subjects with poor to very poor hearing were excluded from reporting normative data on the MMSE, Digit Span and the Rivermead Behavioural Memory Test (RBMT).

### Neuropsychological testing

We encouraged the presence of a close relation during study visits. Close relations were requested to not interrupt during the administration of the cognitive tests. The test battery took approximately 1.5 hours to complete, and we took short breaks whenever subjects showed signs of fatigue. Subjects were encouraged to use all available devices to support their vision and/or hearing. Tests were aborted when sensory problems clearly interfered with test performance. When tests could not be administered or needed to be aborted, we annotated the reasons for interference with test completion: physical, vision or hearing problems, fatigue, or incapable of understanding tasks or instructions.

### Measures

#### Neuropsychological testing

The neuropsychological test battery consisted of fifteen well-known tests that evaluate global cognition, pre-morbid intelligence, attention, language, memory, executive and visuo-spatial functions. The test battery has expanded gradually over the course of the study according to new insights and experience with testing centenarians. Originally, the battery was limited to short screening tests including the MMSE and the CDT and it expanded over time with tests that evaluate specific cognitive domains. Accordingly, not all centenarians were presented with the same test battery or the same number of tests. In total, 77% (n=180) of the centenarians were subjected to a test battery including all 15 tests evaluated here. Tests were administered according to standard procedures, unless stated otherwise.

#### Mini-Mental State Examination

The Dutch version of the MMSE [21, 22] was administered to evaluate global cognitive functioning. The MMSE covers multiple cognitive domains including orientation, memory, attention and/or concentration, working memory, language, visual construction and praxis. The score range for the MMSE is 0 – 30, with higher scores indicating better performance.

#### National Adult Reading Test

We addressed pre-morbid intelligence using the DART, a Dutch version of the National Adult Reading Test [23-25]. Subjects were asked to read out loud fifty words with atypical phonemic pronunciation. We presented these words in a large font-size because of the frequent visual limitations in centenarians.

#### WAIS Digit Span Forward and Backward

We used the Digit Span Forward of the Wechsler Adult Intelligence Scale-III to evaluate attention and/or concentration and the Digit Span Backward to measure working memory [26]. The Forward condition involves subjects to repeat sequences of digits, that increase in length. In the Backward condition, subjects are asked to repeat the increasing digit sequences in reverse order. For both the Digit Span Forward and Backward tests, we present norms for the total raw scores (range 0-16) and the total span of digits (range 0-8), with higher scores indicating better performance.

#### Letter and Category Fluency

The Dutch version of the Controlled Oral Word Association Test (COWAT) [27, 28] was administered to evaluate executive functioning and language. In this Letter Fluency task, subjects were asked to name as many words starting with the letter “D”, “A”, and “T”, each in 1 minute. We also used the Animal Fluency (naming animals in 1 and 2 minutes) to evaluate executive functioning, language and semantic memory [29, 30].

#### Rivermead Behavioural Memory Test

Episodic memory was measured with the story recall subtest of the Dutch version of the RBMT [31-33]. The RBMT contains news articles, which are read out loud to the subjects. The subjects were asked to name all story-items they remembered (immediate recall) and this was repeated after a 15-minute interval (delayed recall). When necessary, the subjects were given a cue for helping them recall the story-line; this cue was not included when calculating the total score. We made two adaptations to the test procedure: 1) two articles were read instead of one in order to improve reliability; 2) during recall all remembered items were scored, whether they belonged to the appropriate story-line or not. The score range is 0-42 for both immediate and delayed recall sections, with higher scores indicating better performance.

#### Visual Association Test

The VAT was developed to detect anterograde amnesia and is based on paired associated learning [34, 35]. Subjects are asked to name two interacting visual items (object or animals, e.g. a hedgehog on a chair), of which one item (hedgehog) needs to be recalled afterwards while the other (chair) is used as a cue. We used naming of the items as an additional measure of language functioning. The score range is 0-12, with higher scores indicating better performance.

#### Trail Making Test

We used TMT A for evaluating processing speed and attention, and TMT B to measure mental flexibility [36]. TMT A requires the subjects to connect 15 dots of numbers in numerical order. In TMT B, subjects were asked to alternate between numbers and letters in numerical and alphabetical sequence (1-A-2-B etc.). For TMT A after 180 seconds, and for TMT B after 300 seconds, we ascertained whether the subjects were still determined to proceed. If this not was the case the test was aborted and scores were extrapolated based on the last finished item (number or letter) and the time spent on the test.

#### BADS Key Search

The Behavioral Assessment of the Dysexecutive Syndrome Test Battery (BADS) [37] was developed to measure executive functioning. Here we used the Key Search subtest, which involves a problem-solving task instructing subjects to think of a strategy to find a lost key. Scores range from 0-16, with higher scores indicating better performance. A time limit was not imposed.

#### VOSP Number location

Visuo-spatial orientation was assessed with the Number Location subtest of the Visual Object and Space Perception Battery (VOSP) [38], in which subjects were asked to visualize and point out a specific number that corresponded with the exact location of a dot. Scores range from 0-10, with higher scores indicating better performance.

#### Clock drawing test

Visuo-spatial construction was measured by the Clock Drawing Test (CDT) [39, 40], in which subjects were instructed to draw a clock with the hands at 10 past 11. Because of poor pen-holding skills or tremor in our subjects, we offered a pre-drawn circle [41]. The scores range from 0 to 5, whereas a score of 0 was given in case of no reasonable representation of a clock, and 5 indicated a perfect clock [40]. In this scoring method, the drawing of the circle itself was not considered.

### Demographic and clinical measures

Education level was determined based on the International Standard Classification of Education 1997 (ISCED) [42]. Independence in performing activities of daily living (ADL) was evaluated with the Dutch observation and/or interview version of the Barthel Index [43-46]. Scores ranged from 0-20, where scores of <15 indicate dependence, and scores ≥15 indicate independence in performing ADL [46]. The 15-item version of the Geriatric Depression scale (GDS) was administered to investigate depressive symptoms [47]. Scores ranged from 0-15: ≤5 indicates no depressive symptoms and >5 suggests the presence of depressive symptoms.

### Data analysis

Normative data were generated by multiple linear regression analyses and by reporting means and percentiles.

The multiple linear regression analyses with age, sex, and education as independent variables and cognitive test scores as dependent variables were performed for the cognitive tests separately. For the regression analyses, TMT scores were log-transformed because of non-normal distributions and inverted, so that higher scores indicated better performance. Because of ceiling effects, the VAT, Number Location and the CDT were not included in the regression analyses. The means and percentiles were reported for each test for the total sample and stratified by education level: low education (post-secondary non-tertiary or lower) and high education (short-cycle tertiary or higher). The cognitive test scores were standardized into z-scores in order to I) correlate the number of tests the centenarians were able to complete with their overall mean z-score and II) to visualize the distribution of the test scores using boxplots. P-values <0.05 were considered significant. Statistical analyses were performed using SPSS 22.0 and R 3.4.2.

## RESULTS

### Demographic and clinical characteristics

The total sample of N=235 had a median age of 100.4 years (range 100-107) at inclusion and included 168 females (71%). The majority (59%) of the centenarians lived independently, 79% were independently mobile and 54% were independent in performing ADL. Most of the centenarians had retained moderate-good vision (77%) and hearing capacities (88%). The majority of the centenarians (62%) had a basic-low education level. Most centenarians (92%) did not show depressive symptoms as measured with the GDS. Clinical and demographic characteristics for this group are summarized in Table 1.

**Table 1.** Demographic and clinical characteristics of the sample (n=235)

| Characteristic |  |
| --- | --- |
| Age, median (IQR)<sup>a</sup> | 100.4 (100.2-102) |
| Age range | (100-107) |
| Sex, N female (%) | 168 (72) |
| Education, N (%) |  |
| High level | 90 (38) |
| Low level | 145 (62) |
| Vision, N (%) |  |
| Good | 153(65) |
| Moderate | 27 (12) |
| Poor | 29 (12) |
| Very poor | 22 (9) |
| Hearing, N (%) |  |
| Good | 134 (57) |
| Moderate | 73 (31) |
| Poor | 21 (9) |
| Very poor | 4 (2) |
| Living situation, N (%) |  |
| Independent without assistance, or in a residence with available service | 138 (59) |
| In a residential care center | 85 (36) |
| In a nursing home | 2 (1) |
| With family | 10 (4) |
| Barthel Index, N (%) |  |
| ≥15 Independent in ADL | 126 (54) |
| <15 Dependent in ADL | 80 (34) |
| GDS >5, depressive symptoms, N (%) | 19 (8) |
| Mobility, N (%) |  |
| Able to walk independently<sup>b</sup> | 185 (79) |
| Able to walk with help of another person | 8 (3) |
| Able to move independently in a wheelchair | 14 (6) |
| Not able to move independently in a wheelchair | 12 (5) |

Values are presented by N and (%), or <sup>a</sup>median (interquartile range). ADL=Activities of Daily Living. GDS=Geriatric Depression Scale. There were 3 missing values for hearing, 4 for vision, 16 for mobility, 45 for the GDS and 29 for ADL independence. <sup>b</sup>=with or without help of a walking stick or walker. High education level=short-cycle tertiary or higher. Low education level=post-secondary non-tertiary or lower.

### Influence of age-related sensory impairments on test incompletion

Across all tests, an average of 79% of the tests was completed (Table 2). While >95% of the subjects completed both fluency tasks, only 45% were able to complete TMT B. Difficulties with vision (41%) and fatigue (22%) were the most common reasons for not being able to complete a test, whereas hearing impairment only rarely complicated test completion (4%). In some cases, not understanding the test and/or test instructions was a reason for incompletion of the Number Location (16%), Key Search (14%), and TMT B (23%). Overall, we found a positive correlation between the number of tests a centenarian was able to complete and the mean z-score across all completed tests (Pearson’s correlation, *r*=.35, *p*<.001), see supplemental data.

**Table 2.** Overview of sample sizes used for generating normative data

| Tests | Tests presented, *N* | Tests completed, *N* (%) | Reasons for test incompleteness, % |  |  |  |  |  | Tests after exclusion for sensory losses, *N* |
| --- | --- | --- | --- | --- | --- | --- | --- | --- | --- |
|  |  |  | No compr. | Fatigue | Hearing | Vision | Several | Other<sup>C</sup> |  |
| MMSE | 235 | 177 (75) | 0 | 0 | 9 | 60 | 12 | 19 | 151<sup>ab</sup> |
| RBMT | 201 | 175 (87) | 0 | 23 | 42 | 0 | 0 | 35 | 167<sup>b</sup> |
| Number Location | 201 | 152 (76) | 16 | 10 | 0 | 63 | 0 | 10 | 142<sup>a</sup> |
| Key Search | 209 | 152 (73) | 14 | 19 | 0 | 47 | 4 | 16 | 138<sup>a</sup> |
| Clock Drawing Test | 234 | 181 (77) | 0 | 13 | 0 | 60 | 6 | 21 | 162<sup>a</sup> |
| Letter Fluency | 226 | 214 (95) | 0 | 50 | 0 | 0 | 0 | 50 | 214 |
| Animal Fluency 1 min | 203 | 196 (97) | 0 | 43 | 0 | 0 | 0 | 57 | 196 |
| Animal Fluency 2 min | 204 | 196 (96) | 0 | 50 | 0 | 0 | 0 | 50 | 196 |
| VAT Memory | 229 | 178 (78) | 0 | 22 | 0 | 57 | 0 | 22 | 156<sup>a</sup> |
| VAT Naming | 206 | 153 (74) | 0 | 19 | 0 | 53 | 0 | 28 | 132<sup>a</sup> |
| Digit Span Forward | 218 | 178 (82) | 0 | 40 | 13 | 0 | 3 | 45 | 163<sup>b</sup> |
| Digit Span Backward | 218 | 180 (83) | 0 | 47 | 13 | 0 | 3 | 37 | 165<sup>b</sup> |
| TMT A | 202 | 133 (66) | 1 | 26 | 0 | 49 | 7 | 16 | 127<sup>a</sup> |
| TMT B | 202 | 91 (45) | 23 | 19 | 0 | 32 | 8 | 19 | 90<sup>a</sup> |
| DART | 228 | 169 (74) | 0 | 24 | 0 | 58 | 2 | 17 | 153<sup>a</sup> |
| Average | 214 | 168 (79) | 6 | 22 | 4 | 41 | 4 | 23 |  |

Columns represent respectively: total number of tests that the centenarians were subjected to, total number of tests that could be completed, reasons for inability to complete tests and total number of tests used for generating normative data after exclusion for sensory losses. MMSE=Mini-Mental State Examination. RBMT=Rivermead Behavioural Memory Test. VAT=Visual Association Test. TMT=Trail Making Test. DART= Dutch Adult Reading Test. No compr.=No comprehension of tests and/or testinstructions. <sup>a</sup>=centenarians with poor to very poor vision were excluded. <sup>b</sup>=centenarians with poor to very poor hearing were excluded. <sup>c</sup>Other includes problems with test equipment, reasons were not reported, physical impairments (tremor or motor), or when there was no time left to finish the whole test battery.

### Normative data and cognitive test performance across centenarians

Per test, an overview of the sample-sizes used for generating normative data is shown in Table 2. Centenarians with poor to very poor vision (21%) and hearing (11%) were excluded for tests for which these faculties were required. We present percentiles and means of all cognitive tests stratified by educational level (Table 3). In the total sample, the mean ±SD score on the MMSE was 25.6 ±3.1 (range 17-30, interquartile-range 24-28). High-educated centenarians scored on average 26.5 ±3.0, while the centenarians who attained lower education levels scored lower (25.2 ±3.1). In addition, multiple linear regression analyses were performed to provide normative data adjusted for sex, age, and education (Table 4). For clinical utility, we also provide the regression formulas for all tests (supplementary material). Correlations between the cognitive test scores are displayed in the supplement. Figure 1 shows the distribution of the performances on each test. Overall, most test scores showed wide distributions, while the VAT, Number Location and the CDT had strong ceiling effects (Table 3).

**Figure 1.**
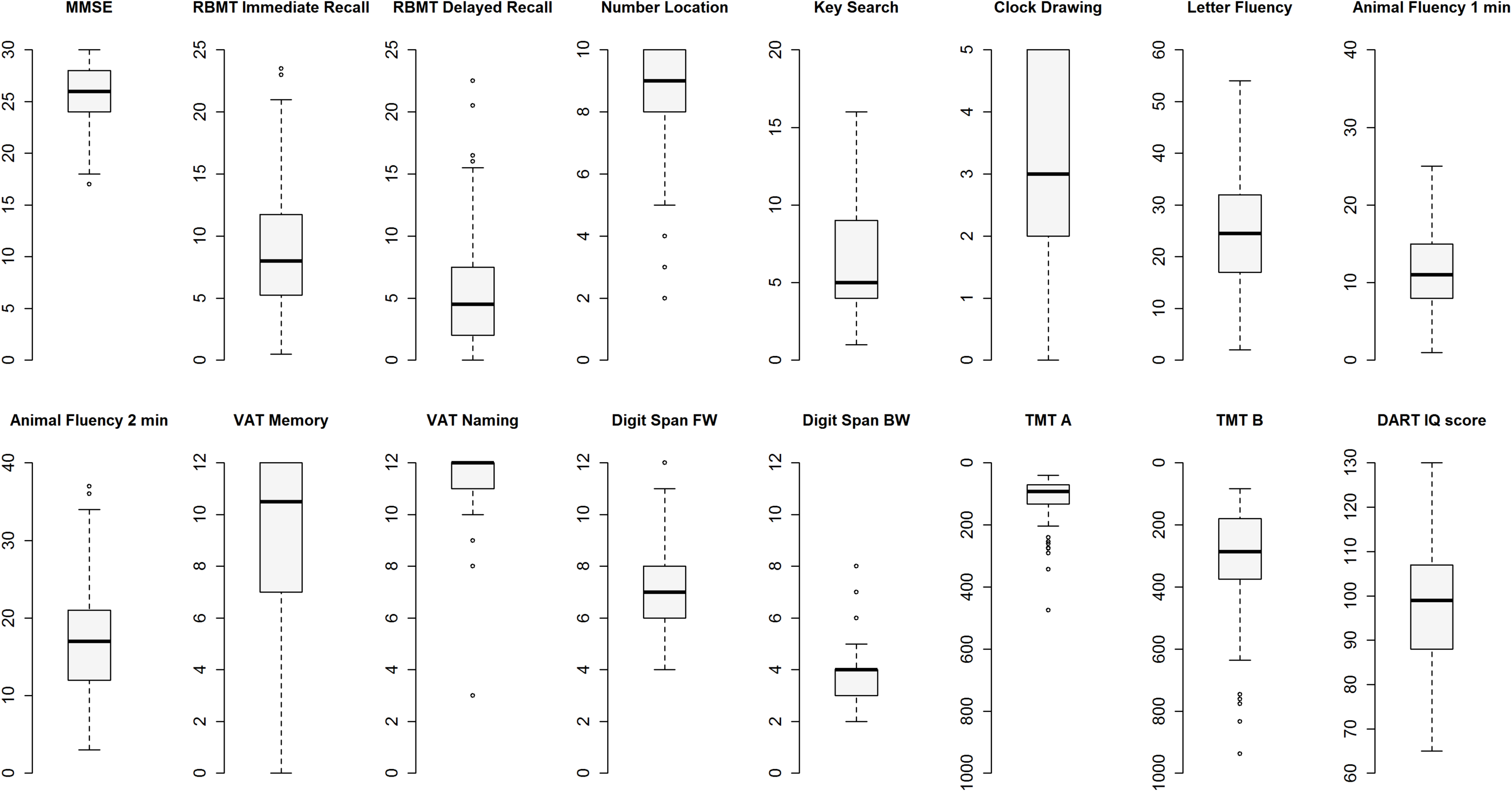
Distribution of neuropsychological test scores. Boxplots represent raw test scores. MMSE=Mini-Mental State Examination. RBMT=Rivermead Behavioural Memory Test. VAT=Visual Association Test. FW=Forward. BW=Backward. TMT=Trail Making Test. DART= Dutch Adult Reading Test.

**Table 3.**
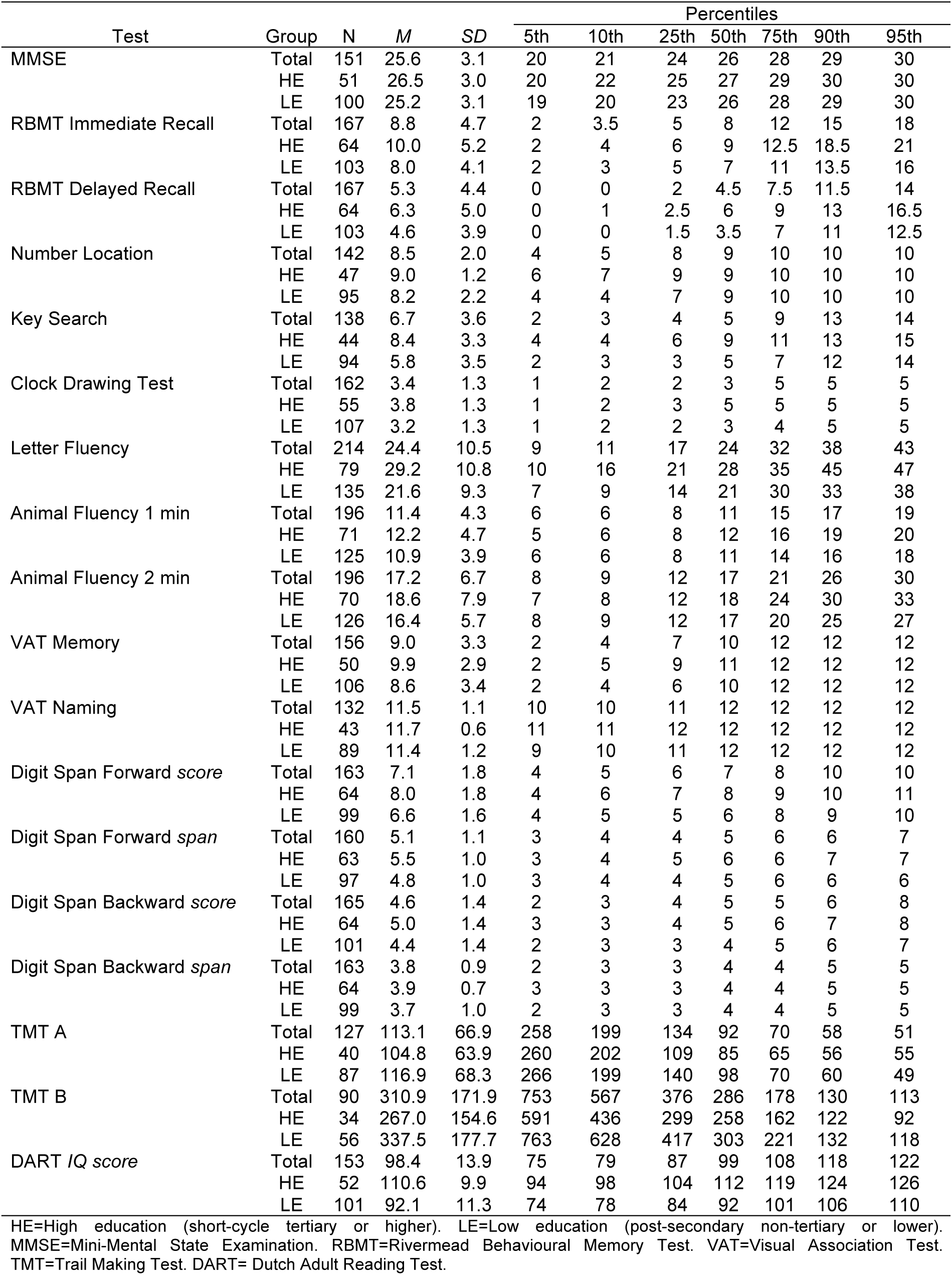
Percentiles and means for cognitive test scores for the total sample and stratified by educational attainment

**Table 4.** Multiple linear regression analyses with sex, age and education as independent variables and cognitive test outcome as dependent variable

| Tests | $R^2$ | Sex | | | Age | | | Education | | |
| --- | --- | --- | --- | --- | --- | --- | --- | --- | --- | --- |
|  |  | *B* | *SE* | *p* | *B* | *SE* | *p* | *B* | *SE* | *p* |
| MMSE | 0.08 | -0.60 | 0.58 | 0.30 | -0.28 | 0.23 | 0.23 | 0.51 | 0.17 | **<.001** |
| RBMT Immediate Recall | 0.05 | -0.64 | 0.79 | 0.42 | -0.10 | 0.31 | 0.73 | 0.61 | 0.24 | **0.01** |
| RBMT Delayed Recall | 0.03 | -0.70 | 0.76 | 0.36 | -0.10 | 0.29 | 0.73 | 0.45 | 0.23 | 0.05 |
| Key Search | 0.16 | -1.71 | 0.65 | **0.01** | -0.04 | 0.26 | 0.86 | 0.75 | 0.19 | **<.001** |
| Letter Fluency | 0.16 | 3.74 | 1.50 | **0.01** | -0.01 | 0.51 | 0.98 | 2.66 | 0.43 | **<.001** |
| Animal Fluency 1 min | 0.05 | 1.24 | 0.67 | 0.06 | -0.17 | 0.23 | 0.46 | 0.54 | 0.20 | **0.01** |
| Animal Fluency 2 min | 0.05 | 1.72 | 1.05 | 0.10 | -0.26 | 0.36 | 0.47 | 0.94 | 0.31 | **<.001** |
| Digit Span Forward *score* | 0.16 | -0.59 | 0.29 | **0.04** | 0.09 | 0.12 | 0.43 | 0.41 | 0.09 | **<.001** |
| Digit Span Forward *span* | 0.11 | -0.15 | 0.18 | 0.40 | 0.03 | 0.07 | 0.64 | 0.23 | 0.05 | **<.001** |
| Digit Span Backward *score* | 0.06 | 0.20 | 0.24 | 0.42 | 0.01 | 0.10 | 0.92 | 0.23 | 0.07 | **<.001** |
| Digit Span Backward *span* | 0.04 | 0.12 | 0.16 | 0.44 | -0.04 | 0.06 | 0.52 | 0.12 | 0.05 | **0.01** |
| TMT A | 0.07 | 0.05 | 0.09 | 0.63 | -0.10 | 0.04 | **0.01** | 0.05 | 0.03 | 0.09 |
| TMT B | 0.07 | 0.23 | 0.12 | 0.06 | -0.05 | 0.05 | 0.38 | 0.06 | 0.03 | 0.08 |

Values are presented as unstandardized Beta (*B*) and standard error (*SE*). MMSE=Mini-Mental State Examination. RBMT=Rivermead Behavioural Memory Test. VAT=Visual Association Test. TMT=Trail Making Test. DART= Dutch Adult Reading Test. TMT scores were log-transformed.

### Association of education, sex, and age with cognitive test performance

The level of educational attainment was positively associated with the performance on all tests when adjusted for age and sex, except the RBMT Delayed Recall, TMT A and B. Sex differences were found for the Digit Span Forward, Key Search and Letter Fluency, with males having higher test scores on the Digit Span Forward (*mean±*SD =7.7 ±1.8 versus females 6.9 ±1.8), and the Key Search (*mean*=8.1 ±4.1 vs *mean*=6.1 ±3.3). Females had higher test scores on the Letter Fluency (*mean*=25.0 ±10.4) than males (*mean*=22.9 ±10.6). On all other tests, males and females performed similarly. Age was only associated with the performance on the TMT A when adjusted for sex and education. The results of these analyses are presented in Table 4.

## DISCUSSION

A large sample of cognitively healthy centenarians was used to generate normative data of fifteen common neuropsychological tests across several cognitive domains, whilst taking vision and hearing impairments into account. We found that test completion was most often complicated by vision impairments and fatigue, and to a lesser extent by hearing impairments or task incomprehension. Almost all cognitive test scores were associated with educational attainment.

### Cognitive test performance in centenarians

Previous studies reported that the variability in cognitive test performance increased with age [11, 12]. Accordingly, most test scores showed wide distributions in the current study, indicating that cognitive test performance was variable within the centenarians. For the VAT, Number Location and the CDT we observed ceiling effects, suggesting that these tests are relatively easy to complete, and might be limited in their ability to capture cognitive performance for this specific population. In line with previous studies, we found that some subjects had difficulty to complete executive functioning tests, supporting the theory that executive functioning is particularly vulnerable to decline in normal ageing [6, 10]. In contrast, the fluency tests were completed by almost all centenarians with varying results and without floor- and ceiling effects, implying that these are suitable tests for application to the oldest-old.

As expected, this selected sample of centenarians scored on average 25.6±3.1 points on the MMSE, which is above the suggested cut-off score used for indicating cognitive impairment in elderly aged 97 years and above [48]. This score is similar to the mean MMSE score of ~24 obtained by American cognitively intact centenarians [15], and markedly higher than the scores of centenarians from population-based studies, which varied from 12.5 to 20 [12-14, 49-51]. More specifically, the test performance of the centenarians on the MMSE, Animal and Letter Fluency, Digit Span Backward, and TMT B, was comparable to scores of the non-demented subjects aged >95 years from the 90+ Study, while performance of the centenarians on Digit Span Forward and TMT A was lower [10]. This could indicate that processing speed and attention may decline in the years between 95 and 100+, while other cognitive domains remain stable. Further, despite selection against vision and hearing impairments in the sample of centenarians, the difference in cognitive test performance might be explained by their increased vulnerability for sensory decline.

### Influence of age-related sensory impairments on test incompletion

Overall, we found an association between the ability to complete tests and test performance, which emphasizes the importance of taking into account factors that interfere with test completion when assessing cognitive functions. Our results suggest that visual impairments, but not hearing impairments, were the most common reason for the inability to complete tests, which is in agreement with previous reports [10]. Therefore, we caution that tests that require vision ability may not be fully applicable in centenarians and that there is room for the design of tests that can be applied independently of these vision impairments. Next to visual impairments, also fatigue commonly led to test incompletion, suggesting that our test battery may have been too extensive for a subset of the centenarians. Therefore, to prevent fatigue from interfering with test results, tests and test batteries for the oldest-old should be kept as short as possible [10, 52].

### Influence of education, sex and age on cognitive test performance

Consistent with previous findings, we found education to be associated with performance on almost all tests, except for delayed recall and the TMT [5, 10]. Accordingly, we expect that having a lower educational attainment is representative of the lower range of the normative data. More specifically, previous studies showed that lower educated elderly scored below cut-off scores on cognitive screening tests, which could cause an overestimation of cognitive impairment in these elderly [53-56]. This might explain that some centenarians, while appearing cognitively healthy during study visits, scored <23 points on the MMSE, or had poor performances on tests on which the majority of the centenarians obtained maximum scores. Therefore, we emphasize that low performance on individual tests should be interpreted in the context of other test scores, taking demographic characteristics and factors of sensory decline into account, to evaluate cognitive impairment in centenarians.

With regard to sex, *population-based* centenarian studies indicated that males consistently performed better on cognitive tests [17, 57], which might reflect the higher dementia prevalence in centenarian females [58, 59]. However, our inclusion criteria of “cognitively healthy centenarians” may introduce a selection bias for only the cognitively healthy males and females, which might explain why we observe no clear sex difference in cognitive test performance. Finally, whereas age is seen as one of the most important predictors for cognitive decline, this was not a strong predictor for cognitive performance in centenarians. We expect that the age range of 100-107, with an interquartile range of 100-101, was too narrow to investigate the effect of age on cognitive test performance.

### Strengths and limitations

In this study we generated normative data specifically tailored for assessing cognitive performance in centenarians, which, given their high risk of having or developing symptoms of cognitive impairment is expected to be variable. The availability of a relatively large sample of cognitively healthy centenarians allowed us to exclude centenarians with sensory losses for the generation of normative data for tests that require vision and hearing ability. Since our test battery expanded during the course of the study, not every test was administered to all subjects. Accordingly, the sample sizes for reporting normative data were unequal and consequently, the order of the tests in the battery could not be taken into account when assessing the influence of fatigue. Further, to select a cognitively healthy population we relied on self-reported cognitive health, which was confirmed by us and by an informant, however, subjects were not screened for dementia according to standard NIA-AA criteria. Nevertheless, we find that the centenarians included in the current study are heterogeneous based on several aspects but based on the MMSE scores, they can be regarded among the highest performing centenarians. Therefore, our cohort of centenarians is appropriate for the generation of normative data to evaluate cognitive function in centenarians.

### Conclusion

This study provides normative data on a broad range of cognitive tests in a large sample of cognitively healthy centenarians, taking hearing and vision impairments into account. The norms established in the current study are useful tools for the detection of dementia and cognitive impairment in centenarians. Our results suggest that, besides vision impairment, also fatigue and education level should be considered when evaluating test results in centenarians.

## Acknowledgements

We are grateful for the collaborative efforts of all participating centenarians and their family members and/or relations. We acknowledge the persons who visited and/or recruited the centenarians. We thank Sven van der Lee, Niccolò Tesi and Wiesje van der Flier who advised on the analyses and presentation of the normative data. This work was funded by Stichting Alzheimer Nederland (WE09.2014-03), Stichting Dioraphte (VSM 14 04 1402) and Stichting VUmc Fonds. The authors declare no conflict of interest.

